# Microbial communities living inside plant leaves or on the leaf surface are differently shaped by environmental cues

**DOI:** 10.1101/2023.12.17.572047

**Authors:** Maryam Mahmoudi, Juliana Almario, Katrina Lutap, Kay Nieselt, Eric Kemen

## Abstract

Leaf-associated microbial communities can promote plant health and resistance to biotic and abiotic stresses. However, the importance of environmental cues in the assembly of the leaf endo- and epi-microbiota remains elusive. Here we aimed to investigate the impact of seasonal environmental variations, on the establishment of the leaf microbiome, focusing on long-term changes (five years) in bacterial, fungal, and non-fungal eukaryotic communities colonizing the surface and endosphere of six wild *Arabidopsis thaliana* populations. While leaf-microbial communities were found to be highly stochastic, the leaf niche had a predominant importance with endophytic microbial communities consistently exhibiting a lower diversity and variability. Furthermore, our analyses revealed that among environmental factors, radiation and humidity-related factors are the most important drivers of diversity patterns in the leaf, albeit with stronger effects on epiphytic communities. Using linear models, we further identified 30 important genera whose relative abundance in leaf compartments could be modeled from environmental variables, suggesting specific niche preferences for these taxa. With the hypothesis that these environmental factors could impact interactions within microbial communities, we analyzed the seasonal patterns of microbial interaction networks across leaf compartments. We showed that epiphytic networks are more complex than endophytic, and that the complexity and connectivity of these networks are partially correlated with the mentioned environmental cues. Our results indicate that humidity and solar radiation function as major environmental cues shaping the phyllosphere microbiome at the micro-scale (leaf compartment) and macro-scale (site). These findings could have practical implications for selecting and developing field-adapted microbes in the face of, and for predicting microbial invasions in response to global change.

## Introduction

Leaves are colonized by various microbes including bacteria, fungi, oomycetes, and protists [1]. This leaf microbiota can play a beneficial role in protecting plants against biotic and abiotic stressors, thus ultimately promoting plant growth and fitness [2–6]. Within the microcosm of the leaf, distinct compartments emerge, primarily characterized as epiphytic (surface) and endophytic (internal tissues). Despite their proximity, these zones have different characteristics. The leaf surface is covered by a hydrophobic cuticle layer that prevents water loss from the leaf surface. This environment comprises components like wax and cutin [7], along with trichomes, which can protect against UV light and mediate leaf temperature [8]. Whereas, within the leaf interior, a vast area known as the apoplast that facilitates gas and water exchanges for photosynthesis. This environment, with its higher humidity, is also subjected to microbial colonization [9]. In addition, the apoplast can be subject to pH fluctuations in response to biotic stresses such as pathogen attack and abiotic factors such as salinity or drought [10]. Due to their different characteristics, these two niches may favor certain microbes and make it difficult for others to survive. In this context, it is not well known how environmental cues differentially shape the microbial communities occupying these two niches.

Environmental factors (e.g., light and humidity) significantly affect all microbial communities in different ecosystems and the phyllosphere [11, 12]. Light, as a fundamental element of living organisms, is essential not only for plant photosynthesis and growth but it also mediates plant-microbe interactions. The presence of photoreceptor proteins in microorganisms enables them to detect light to use for adhesion to host tissues for colonization, and for DNA repair [13]. In particular, UV light can enhance plant defense mechanisms by stimulating the production of defense-related compounds such as salicylic acid and jasmonic acid, which strengthen plants against pathogens [14]. In addition, the availability of water and nutrients is critical for plant health and ecosystem balance [15]. Precipitation has been shown to play a significant role as a primary driver in shaping fungal communities and facilitating the spread of fungal plant pathogens via rain droplets [16, 17]. Accordingly, precipitation has a significant effect on the composition of soil microbial communities [18]. Although some research has investigated the effects of environmental factors on leaf microbiomes, there remains a lack of studies focusing on the effects of such factors on different microbial communities within leaf compartments.

Microbes often interact with each other through various relationship types, such as mutualism or antagonism, and can develop complex plant-associated communities that can change throughout the growing season of the host [19, 20]. The use of microbial interaction network analysis has been useful in understanding the variability and stability of these communities under changing environments [21–23]. For example, the complexity of microbial networks has been linked to community stability, as seen in a long-term study of grassland soil microbiome, showing that warming increases the complexity of microbial network (e.g., size and connectivity) [24]. Environmental stresses (decreasing water availability, nutrients, and vegetation) have been shown to have impact the stability of microbial communities by decreasing richness and altering the ratio of positive and negative interactions [25]. However, how plant-associated microbial communities respond to changing environmental conditions has rarely been studied using multi-kingdom microbial interaction networks.

Microbiota associated with plants and animals undergo seasonal fluctuations shaped by environmental cues and perturbations [26, 27]. In plants, research has shown that the microbiome tends to become more tissue-specific throughout the host developmental stages [28], but these communities remain highly stochastic. For example, tracking the leaf microbiome of *Arabidopsis thaliana* during its growing season from November to March in common garden experiments revealed overall high variability with some conserved patterns [29]. These conserved patterns were characterized by identifying more persistent microbes known as core microbes. Among these core microbes, some members of plant pathogens, such as *Peronosporales*, increased throughout the growing season, reaching maximum values in March which aligns with the disease dynamics of downy mildew in *Brassicaceae*, known to be favored by cold, wet weather [29]. However, despite the recognized importance of longitudinal microbiome data, few studies have used environmental data to explain observed temporal dynamics in the plant-associated microbiota.

The objective of this research was to link temporal changes in the leaf microbiome of natural *Arabidopsis thaliana* populations, with naturally occurring environmental factors, in a long-term study. We hypothesized that leaf-associated microbiomes occupying the epiphytic or the endophytic compartment would respond differently to environmental cues. Using amplicon sequencing, we tracked leaf microbial communities (bacteria, fungi, and non-fungal eukaryotes) during fall and spring seasons, over five consecutive years. Our results revealed that while many environmental factors shaped these communities, the leaf niche emerged as the most important factor. Endo- and epiphytic microbial communities exhibited distinct responses to environmental cues, with radiation- and humidity-related factors appearing to have a greater influence on the diversity and structure of these communities. We further identified 30 microbial taxa showing distinct responses to certain environmental cues, suggesting some level of niche preference among leaf-colonizing taxa. By examining microbial interaction networks between epiphytic and endophytic communities, we further found that community cohesion, a measurement of connectivity, could be correlated with specific environmental factors suggesting that certain environmental cues can drive community stability.

## Results

### Leaf epiphytic and endophytic microbial communities differ in diversity and structure

With the aim to study the impact of environmental cues on the temporal dynamics of the leaf endo- and epi-phytic microbiota, we collected samples from six locations (sites) with stable *A. thaliana* populations in the proximity to Tübingen (south Germany)(Fig. 1A) [30] over two seasons (fall and spring), and five years (Fig. 1C; see Table. S1). We then correlated changes in the leaf microbiota with environmental variables measured locally (14 environmental factors with monthly resolution, Table. S2). Fall sampling covered the early growth phase of *A. thaliana* while Spring sampling covered the early reproductive stage. From each sample, we recovered epiphytic and endophytic microbial communities, extracted genomic DNA, and performed bacterial 16S rRNA, fungal ITS2, and eukaryotic 18S rRNA amplicon sequencing (Fig. 1D), as described in [30]. In the analysis of the 18S eukaryotic data, all microbes that belonged to the kingdom fungi were excluded. We refer to this data as non-fungal eukaryotes (NFEuk).

**Figure 1.**
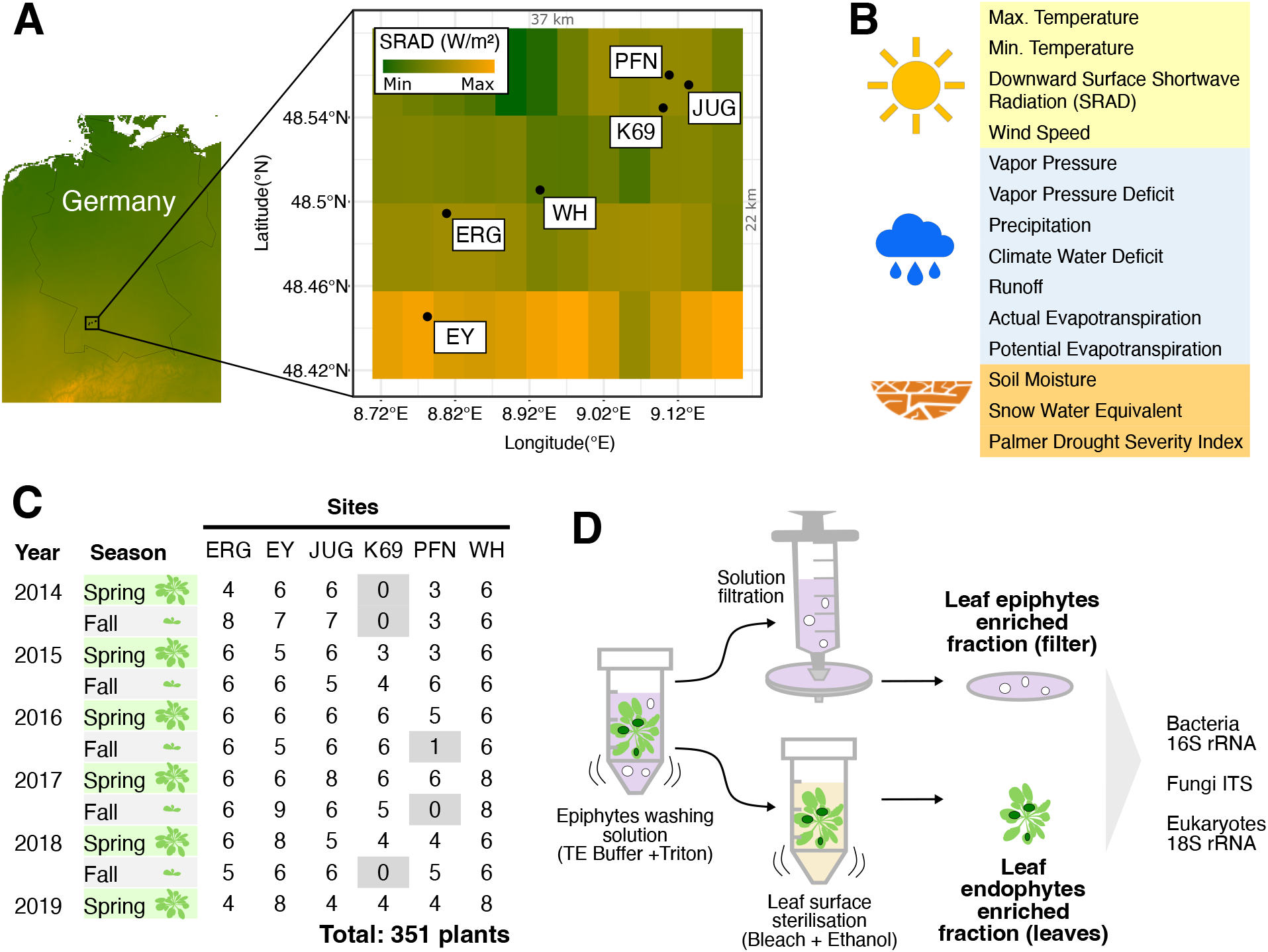
Microbial community collection in natural *A. thaliana* populations over time. **(A)** Map showing the six sampling locations of natural *A. thaliana* in southern Germany near Tübingen [30]. The heatmap on the map represents average variation in solar radiation of sampling locations (downward surface shortwave radiation (srad)). **(B)** Environmental variables (14; Table. S2) used in this study were obtained from the TerraClimate database [47]. **(C)** Plants (n = 351) were collected in the fall and spring of five consecutive years (starting spring 2014, ending spring 2019, 11 time points). **(D)** Leaf epiphytic and endophytic fractions collected from each sampled rosette (Table. S1). Microbiome analysis was conducted via Illumina-based amplicon sequencing (Miseq 2×300 bases). Taxonomic markers included the bacterial 16S rRNA V5-V7 region, fungal ITS2, and 18S rRNA V9 region of eukaryotes.

To investigate the effect of the “compartment” (endophytic vs epiphytic fractions; Fig. 1), the “site” and the “season” on leaf-associated microbial communities, we conducted multiple diversity analyses. Permutational multi-variate analysis of variance (PERMANOVA) results show that the leaf “compartment” emerges as the primary driver of the structure of microbial communities (Fig. 2A). In particular, it exerts a major influence on the structure of bacterial (8.4% explained variance) and non-fungal eukaryotic communities (11.8%). Fungal communities appeared much less constrained by the leaf compartment (2.3% explained variance) and were more influenced by the sampling site (5.7%). These analyses further revealed a marginal effect of the “season” (explaining 0.8% to 3% of the variance). Accordingly, non-metric multidimensional scaling (NMDS) plots showed a clear separation between epiphytic and endophytic samples for bacterial and non-fungal communities, while fungal communities exhibited the smallest separation (Fig. 2B).

**Figure 2.**
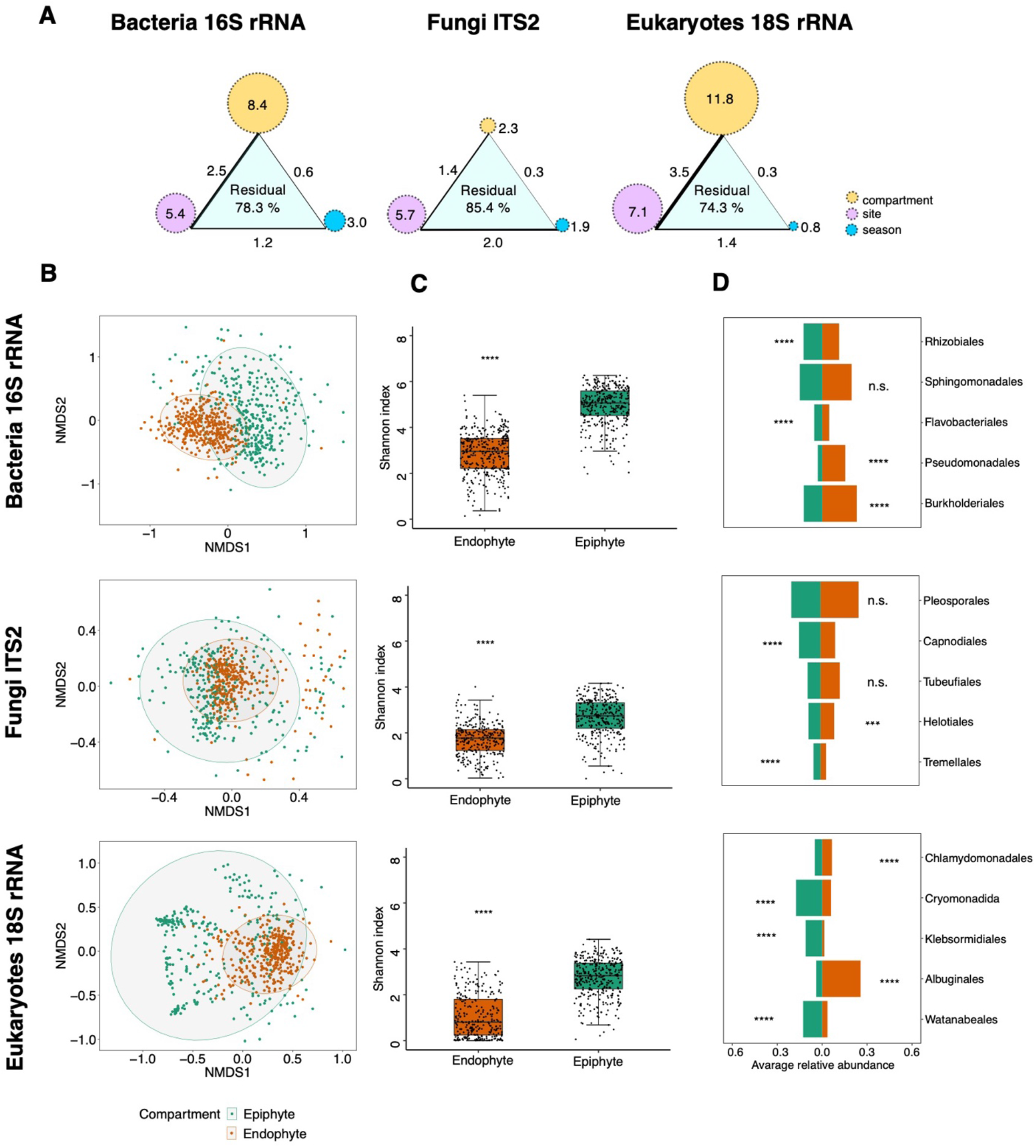
Multivariate analysis on factors structuring leaf communities. **(A)** A PERMANOVA analysis on Bray-Curtis dissimilarities. Circles depict the percentage of variance explained by factors ‘compartment’, ‘site’, and ‘season’; connecting lines depict the percentage of variance explained by interactions between factors, and the thickness of lines shows the strength of explained variation. Only significant effects are shown (permutations 10,000, *P* ≤ 0.05, explanatory categorical variables: compartment x site x season). **(B)** Non-metric multidimensional scaling ordination (NMDS) analysis of epiphytic and endophytic samples measured by Bray-Curtis dissimilarities in bacterial, fungal, and non-fungal eukaryotes. **(C)** Alpha-diversity measures (Shannon’s H index) of epiphyte and endophyte samples. The box plots display individual samples as dots. **(D)** Average relative abundances of the top five most abundant microbial orders in epiphytic and endophytic samples. Asterisks indicate significant differences based on Wilcoxon’s test: n.s. (*P >* 0.05), * (*P* ≤ 0.05), ** (*P* ≤ 0.01), *** (*P* ≤ 0.001), and **** (*P* ≤ 0.0001).

Alpha-diversity measures (Shannon’s index, related to the number of taxa in the community) show that leaf endophytic communities were 1.6 to 2.6 times less diverse than their epiphytic counterparts (Fig. 2C). Additionally, the diversity of the endophytic communities appeared less variable between seasons, than that of epiphytic communities which showed significant changes between seasons (Fig. S1A). A notable exception were endophytic bacterial communities which were significantly more diverse in the fall than in the spring (Wilcoxon’s test, *P <* 0.0001) (Fig. S1A).

Differences in epiphytic and endophytic communities were associated with the enrichment of major microbial orders (overall most abundant orders; Fig. 2D). Among bacteria, *Rhizobiales* and *Flavobacteriales* were more abundant among epiphytes (1.1 times and 1.2 times, respectively), while *Burkholderiales* and *Pseudomonadales* were more abundant among endophytes (1.9 and 5 times more, respectively). Among fungi, *Tremellales* basidiomycetes and *Capnodiales* ascomycetes were more abundant in the epiphytic fraction (1.2 times and 1.4 times, respectively), while ascomycetes from the *Helotiales* were enriched in the endophytic fraction (1.1 times). In addition, non-fungal eukaryotic orders enriched in the epiphytic compartment included green algae *Watanabeales* and *Klebsormidiales*, as well as the cercozoan *Cryomonadida* (3.5 times, 7.8 times and 2.9 times, respectively). Surprisingly, green algae from the *Chlamydomonadales* were 1.3 times more abundant in endophytes. Finally, *Albuginales*, known to harbor the plant biotrophic pathogen *Albugo*, were 6.4 times more abundant among endophytes. These results illustrate the extent of preference that major leaf-associated microbes have for either the epiphytic niche or the endophytic niche.

### Endophytic and epiphytic microbial communities respond differently to environmental cues

We hypothesized that the major differences observed between endo- and epiphytic communities are partially explained by the fact that these communities respond differently to major environmental cues. To test this hypothesis, we evaluated the effect of 14 selected environmental factors on community structure (PERMANOVA on Bray-Curtis dissimilarities) and alpha-diversity (correlation of environmental factors with communities alpha-diversity), in each of these niches. The fourteen environmental variables selected (Fig. 1B and Table. S2) showed variability across seasons, years and/or sampling sites (Fig. S2). While all environmental factors significantly impacted the structure of leaf-associated microbial communities, marginal effects were observed for most of the factors with very low percentages of variance explained (0.5 to 1.9, *P <* 0.05) (Fig. 3A, Table. S3). Notably, bacterial communities were more affected by solar radiation and humidity-associated factors like vapor pressure, precipitation and evapotranspiration (actual and potential) than micro-eukaryotic communities (fungal and non-fungal) (Fig. 3A). More pronounced effects were found when considering the correlations between these environmental factors and per−sample microbial alpha-diversity. While factors associated with temperature (minimum and maximum) and high humidity (vapor pressure, precipitation, and soil moisture) had overall positive effects on microbial alpha-diversity, solar radiation had an overall negative effect. A notable exception was that solar radiation was positively correlated to higher fungal diversity, specifically on the leaf surface (epiphytic fraction) (Fig. 3B). This suggests that increased radiation levels may stimulate the growth or proliferation of certain fungal species adapted to thrive under such conditions.

**Figure 3.**
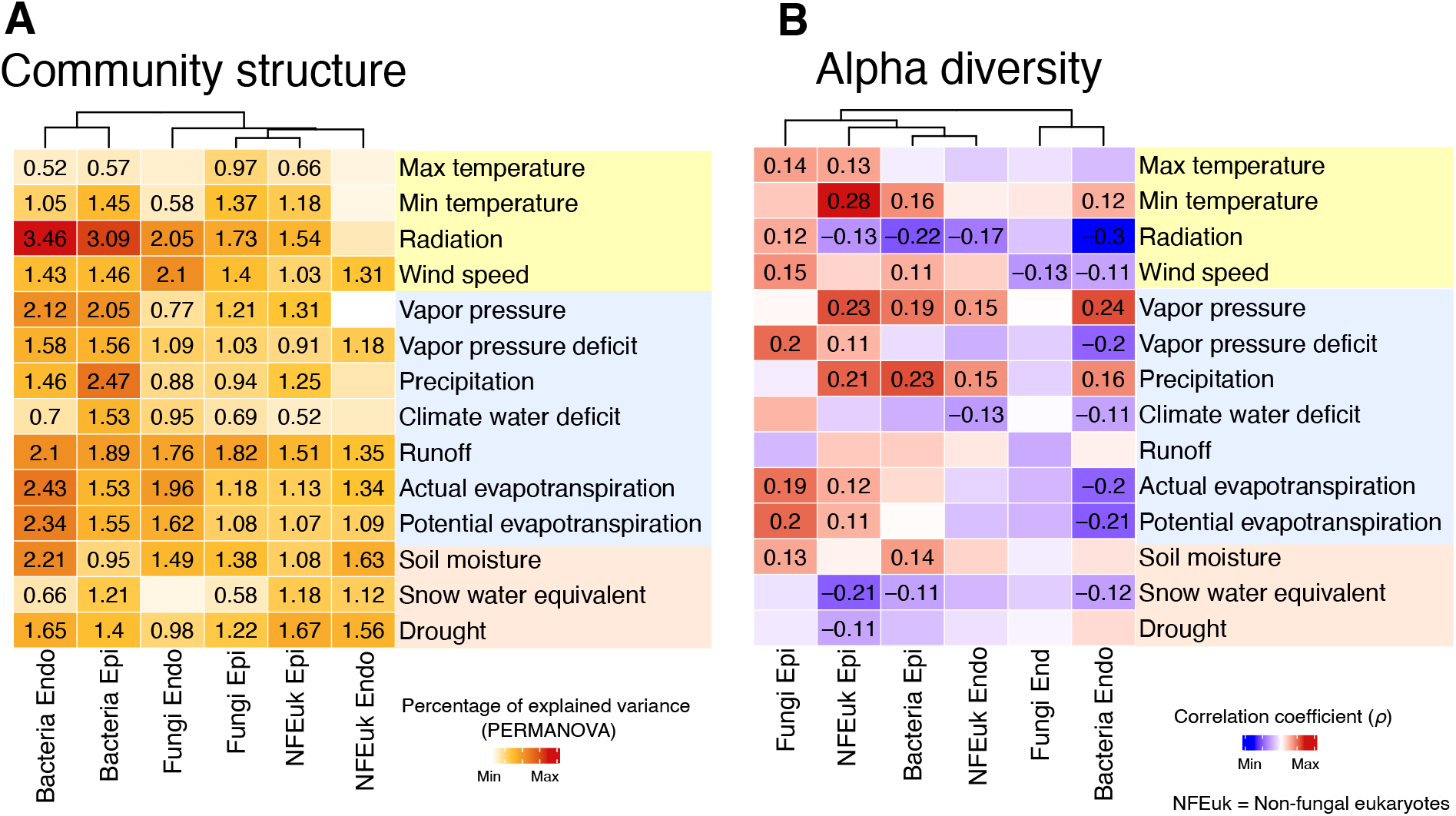
Effects of environmental factors on microbial community structure and alpha diversity. **(A)** Heatmap displaying explained variance in PERMANOVA models (*R*^2^ value; BrayCurtis dissimilarities) indicating the influence of individual environmental factors on microbial community structure, within each leaf compartment. **(B)** Heatmap showing Spearman correlation coefficients (*ρ*) between alpha-diversity (Shannon’s H index) and environmental factors, in each compartment. Only significant results are displayed (*P <* 0.05).

Interestingly, certain factors had stronger effects on one niche or one microbial group. For example, in comparison to their epiphytic counterparts, the alpha-diversity of endophytic micro-eukaryotic communities (fungal and non-fungal), showed overall fewer significant correlations with the analyzed environmental factors (5 vs 17 significant correlations). This aligns with the previous observation that micro-eukaryotic alpha diversity is less variable inside the leaf (endophytic) than on the leaf surface (Fig. 2C and Fig. S1), suggesting that these communities are more resistant and/or resilient to environmental perturbations. Interestingly, these trends did not hold for bacterial alpha-diversity which correlated with several factors both for endophytic and epiphytic communities (10 vs 7 significant correlations). Some factors associated with water loss from the plant (wind speed and actual/potential evapotranspiration) had contrasting effects on microbial diversity depending on the niche considered, with negative effects on endophytic diversity (fungal and/or bacterial communities) and positive effects on epiphytic diversity (bacterial and/or micro-eukaryotic communities). Taken together these results suggest that environmental factors influence microbial communities differently depending on their habitat and their broad phylogenetic group.

### The abundance of specific taxa in different leaf compartments can be inferred from certain environmental data

Further analyses aimed to assess the impact of key environmental factors on the relative abundance of major microbial genera. To this end we used a GLM approach (generalized linear models) to assess the response of selected genera, in each leaf compartment. These analyses revealed significant effects for at least one environmental factoron the relative abundance of most taxa: 91% of the bacterial genera, 85% of the fungal genera and 86% of the non-fungal genera (FDR-corrected *P <* 0.05; Table. S4). When examining the 30 most responsive genera shared between epiphytic and endophytic compartments (those with the highest average coefficient values in the GLMs)), we found that they were mainly impacted by precipitation, soil moisture, maximum temperature, drought, radiation, and vapor pressure. Yet, no single factor was significant for all these taxa (Fig. 4). In this subset, the relative abundance of the considered bacterial genera on the leaf surface (epiphytes) was more often impacted by the considered factors than their endophytic counterparts, but the effects were marginal (coefficients below 0.01). Overall, radiation had a positive effect on the relative abundance of most of these taxa in the epiphytic compartment, while factors associated to high humidity (precipitation, vapor pressure and potential evapotranspiration) had mixed results with negative effects on *Sphingomonas* and positive effects on *Flavobacterium*. For endophytic bacteria, the strongest effects were observed for *Pseudomonas* relative abundance which increased with higher radiation and lower humidity (lower soil moisture and lower potential evapotranspiration), and *Sphingomonas* relative abundance which increased with higher humidity (higher soil moisture and lower wind speed). These differential responses are probably associated with different niche preferences for these taxa.

**Figure 4.**
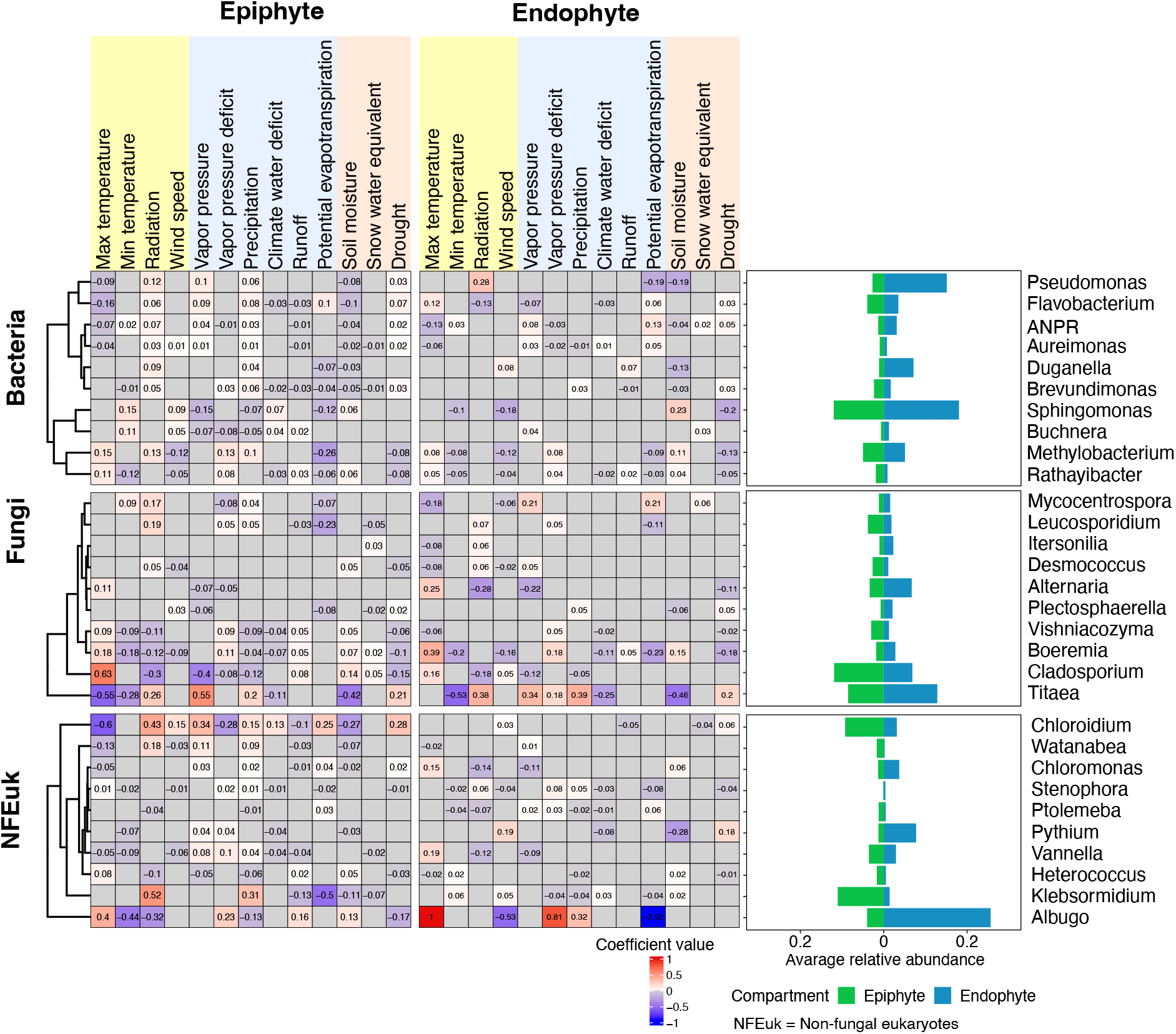
Association between environmental factors and relative abundance of microbial genera among epiphytic and endophytic compartments. **(A)** The values of the heatmaps show coefficient values of environmental factors in modeling the relative abundance of microbial genera using GLM. Negative values indicate genera that decrease with these environmental factors, while positive values indicate genera that increase. Only significant relations (*P <* 0.05, FDR-corrected) are displayed. (B) The histograms display the average relative abundances of selected microbial genera in each compartment. **(B)** Average relative abundance of selected genera present in epiphytic and endophytic compartments, and highly influenced by environmental factors (highest average coefficient values).

Like bacteria, the relative abundance of major fungal taxa on the leaf surface (epiphytes) was more frequently influenced by the considered factors than their endophytic counterparts. However, the effects observed were mostly marginal (coefficients below 0.01). Notably, radiation and precipitation yielded mixed results. High radiation and humidity (precipitation) negatively impacted the abundance of fungal taxa *Cladosporium, Boeremia*, and *Vishniacozyma*, while increasing the abundance of *Titaea* (*Tetracladium*), a *Helotiales* fungus which is known to thrive in water environments [31].

Similarly, the relative abundance of major non-fungal eukaryotic taxa on the leaf surface (epiphytes) was more responsive to environmental factors than their endophytic counterparts, albeit with mostly marginal effects (coefficients below 0.01). Overall, temperature and precipitation had a positive impact on the relative abundance of most taxa in the epiphytic compartment. Specifically, precipitation increased the relative abundance of green algae *Chloridium* and *Klebsormidium* in the leaf surface (epiphytic), which is in line with the fact that these organisms proliferate in light-exposed high-humidity environments. The most striking results were observed with the pathogenic biotroph *Albugo* (oomycete) whose abundance inside the leaf was negatively impacted by high humidity indicators (high potential evapotranspiration, low wind speed) and promoted by high maximum temperatures, suggesting this pathogen invades the leaf under dry heat conditions.

### Microbial networks and community cohesion are driven by major environmental cues

We conducted a microbial network analysis to explore changes in the interactions among microbes in the epiphytic and endophytic compartment, aiming to assess the impact of environmental factors on the connectivity of these communities. Microbial networks were constructed for each of the eleven sampling times, and a comparative examination was carried out between the epiphytic and endophytic networks (Fig. 5A). It is worth noting that the epiphytic network had a greater complexity than the endophytic network with a larger number of nodes (OTUs) and edges (correlations between taxa). On average, the epiphytic compartment contained 15.1 times more nodes and 79.7 times more edges (559 nodes and 3348 edges) than the endophytic compartment (37 nodes and 42 edges).

**Figure 5.**
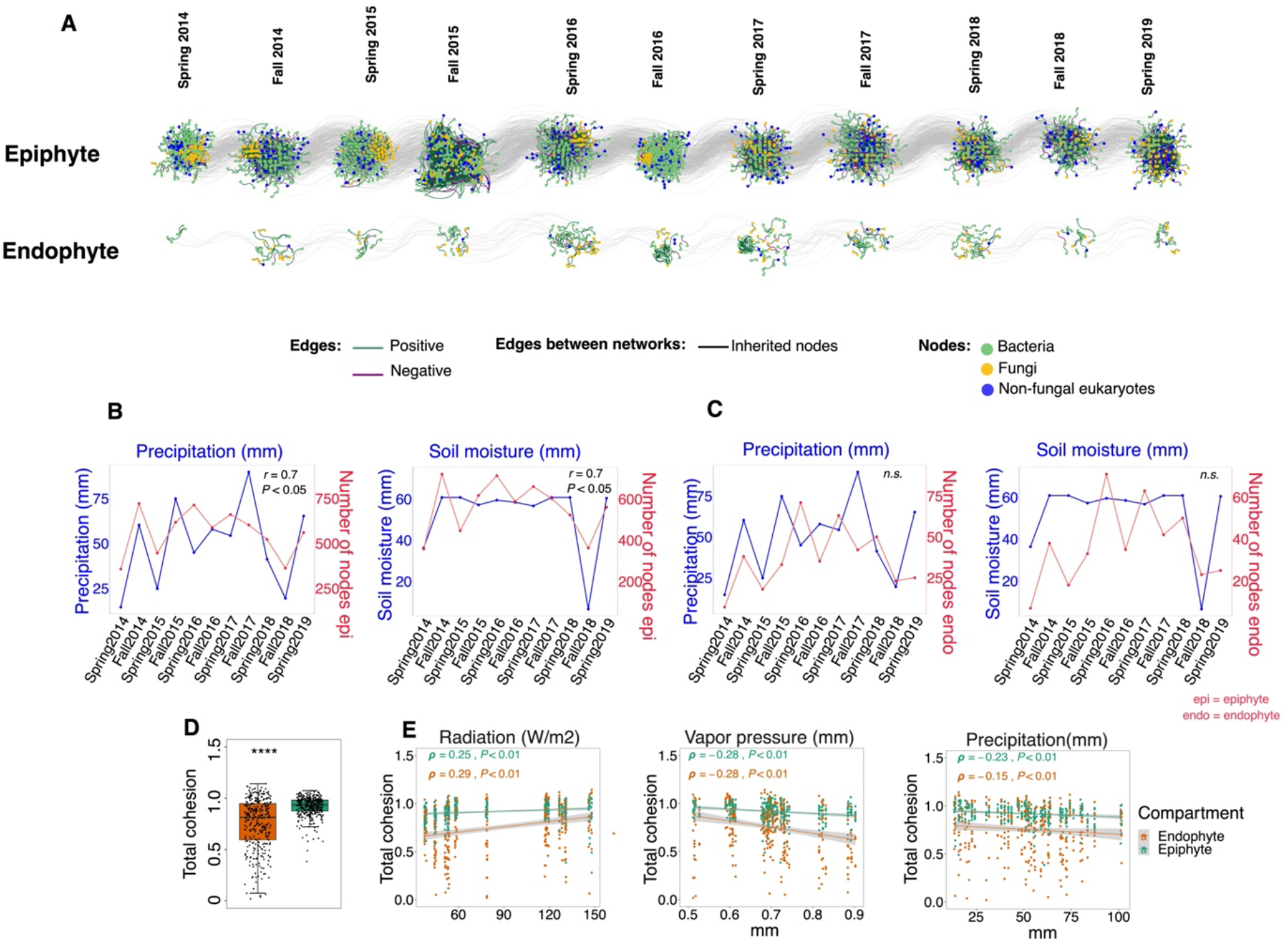
Correlating microbial network complexity and environmental factors. **(A)** Data from each time point was used to reconstruct co-abundance networks for epiphytic and endophytic compartments. The nodes (dots) represent OTUs, and the edges (colored lines) depict potential positive and negative interactions between OTUs (connections). Gray lines (connecting the networks) show nodes conserved in networks from one-time point to the next (inherited nodes). **(B-C)** Correlation between the number of nodes and monthly precipitation and soil moisture in the epiphytic and endophytic compartments across time points (Pearson correlation coefficient *r, P <* 0.05). **(D)** Total cohesion (sum of positive cohesion and the absolute value of negative cohesion) in epiphytic and endophytic samples. **(E)** Correlations between total cohesion and solar radiation, vapor pressure or precipitation, across leaf compartments (Spearman correlation coefficient *ρ, P <* 0.05). The grey lines indicate 95% confidence intervals. Individual samples are represented by dots and colored by compartments.

Further analyses were conducted to investigate the potential correlation between the complexity of microbial interaction networks (number of nodes and connectivity) and environmental factors. The findings revealed that from the 14 factors analyzed only two factors: precipitation and soil moisture, significantly correlated with the number of nodes in epiphytic networks (*r* = 0.7, *P <* 0.05) (Fig. 5B), while no significant correlations could be drawn for endophytic networks (*P* ≥ 0.05) (Fig. 5C, Table. S5).

We further investigated whether the connectivity of microbial communities, which considers the strength of positive and negative interactions, could be explained by these environmental factors. To this end, we computed a community cohesion metric previously proposed [32]. Our analysis unveiled that epiphytic communities exhibited significantly higher cohesion levels compared to endophytic communities (*P <* 0.001) (Fig. 5D). Among environmental factors shaping community cohesion (Fig. S3), radiation exhibited the highest positive effect both for epiphytic and endophytic networks, along with factors associated with low humidity such as low vapor pressure and low precipitation (Fig. 5E). Taken together these results suggest that higher humidity is associated with bigger (more nodes) microbial networks on leaf surfaces, while increased solar radiation and low humidity are associated with overall more connected networks (higher cohesion).

## Discussion

The phyllosphere is a system directly exposed to various environmental factors such as light and humidity. All these factors can cause significant perturbations to the microbiome of the leaf [29, 33]. This leads to a fundamental question: to what extent do seasonal environmental factors determine the different dynamics of epiphytic and endophytic microbial communities? To address this fundamental ecological question, we have conducted a comprehensive, five-year investigation of the leaf microbiome of *Arabidopsis thaliana* from natural populations in six different geographical locations [30] (Fig. 1). Our results highlight the significant influence of plant compartment, site location, and sampling season on shaping microbial communities, elucidating 14.6-25.7% of their variability, including bacteria, fungi, and non-fungal eukaryotes (Fig. 2), consistent with results from previous studies [30, 34–39].

In our analysis, we identified the leaf compartment as the primary factor driving the variation in the bacteria and non-fungal eukaryotes, resulting in a lower level of alpha diversity among the endophytes (Fig. 2). This finding supports the hypothesis that the diversity gap might result from different conditions within these niches. Endophytic microbiomes face obstacles such as apoplastic acidity and oxidative stress [40], as well as nutrient deficiencies [41]. These are likely to affect their diversity patterns. These challenges may therefore be responsible for the observed lower endophyte diversity. Conversely, the observed higher diversity among epiphytes suggests that the leaf surface, which is more exposed to environmental elements, provides more favorable conditions for microbial proliferation than the protected environment within the plant. In addition, it’s important to recognize the potential influence of environmental factors on microbial communities distributed within these compartments.

We found that solar radiation correlated negatively with microbial alpha diversity (Fig. 3). This effect could occur directly by damaging microbial DNA, especially on the leaf surface. Alternatively, it could affect diversity indirectly by promoting the production of reactive oxygen species (ROS) that inhibit the growth and diversity of sensitive species. Long-term low-dose ionizing radiation has been shown to affect soil microbial communities by inhibiting predatory or parasitic fungi [42]. For example, we found a reduction in the abundance of the fungus *Cladosporium* in response to solar radiation. While *Cladosporium* species are melanized filamentous fungi, and melanin can typically protect them from UV radiation, our results contradicted this expectation [43]. Interestingly, we observed an increase in endophytic *Pseudomonas* species favored by solar radiation, possibly due to their pigment-producing abilities. This suggests that bacteria in the plant microbiome may use pigments as a protective mechanism against ROS, which is particularly important under high UV radiation or intense light [44, 45]. While analyzing microbial interaction networks, solar radiation emerged as an important factor positively correlated with their cohesion. This suggests that radiation may increase the strength of interactions. This may represent a survival strategy whereby microbes form stronger bonds or dependencies to cope with the environmental stress imposed by radiation exposure, thereby increasing resilience to external perturbations.

Humidity-related factors, such as precipitation and vapor pressure, emerged as significant contributors to higher microbial alpha-diversity (Fig. 3). This was expected as humidity influences microbial diversity by modifying substrate diffusion [46] and facilitating microbial dispersal via rain, as demonstrated for fungal diversity [16, 17]. However, we found negative correlations between high humidity parameters (precipitation and vapor pressure), with microbial network cohesion. This suggests that under conditions such as precipitation, microbes may choose strategies such as adhesion over motility, potentially reducing the connectivity of microbial interactions. In addition, intense precipitation may physically disrupt microbial habitats and structures, such as biofilms or microbial aggregates, leading to temporary disintegration of microbial networks and reduced cohesion. Alternatively, during periods of high precipitation or humidity, microbial communities may allocate more resources to survival mechanisms such as biofilm formation or stress response pathways rather than investing in microbial interactions.

## Conclusions

Our study conducted a comprehensive analysis of leaf microbiomes over an extended period, revealing the leaf compartment as the primary determinant shaping microbial communities. In addition, we highlighted the critical role of environmental cues in shaping the diversity, composition, and interactions among microbes within leaf compartments. In particular, we identified specific microbial communities that respond to these environmental cues. By using cohesion as a metric to quantify microbial community connectivity [32], we illuminated how external environmental factors can alter internal microbial interactions. Our study introduces a novel approach for investigating temporal community dynamics in natural, host-associated microbiomes. In addition, our findings hold promise for advancing the modeling and prediction of microbial community dynamics over time using insights into environmental influences. Understanding these processes could potentially guide efforts to direct microbial communities toward desired states, with valuable implications for ecosystem management and sustainability.

## Method

### Collection of *Arabidopsis thaliana* samples and environmental data

Wild *Arabidopsis thaliana* samples were collected from six sites near Tübingen. In the fall and spring over five years (2014-2019, 11 time points, Table. S1). Epiphytic and endophytic microorganisms were collected from each sample, as described in Agler *et al*. [30]. In brief, rosettes were washed gently with water for 30 sec, then in 5 ml of epiphyte wash solution (0.1% Triton X-100 in 1x TE buffer) for 1 min. Epiphytic microorganisms were collected by filtering the solution through a 0.2 um nitrocellulose membrane filter (Whatman, Piscataway, NJ, USA). The filter was placed in a screw-cap tube and frozen in dry ice. For collecting endophytic fractions, the rosette was then surface-sterilized by washing with 80% ethanol for 15 sec, followed by 2% bleach (sodium hypochlorite) for 30 sec. Rosettes were rinsed thrice with sterile autoclaved water for 10 sec, before placing them in a screw-cap tube and freezing them on dry ice. Phenol-chloroform-based DNA extraction was performed according to a custom protocol as described in Agler *et al*. [30]. The extracted DNA was used for two-step PCR amplification of the V5-V7 region of bacterial 16S rRNA (primers 799F/1192R), the fungal ITS2 region (primers fITS7/ITS4), and V9 region of eukaryotic 18S rRNA (primers F1422/R1797) (Table. S6). Blocking oligos were used to reduce amplification of plant DNA (Table. S6). Purified PCR products were pooled in equimolar amounts before sequencing in Illumina MiSeq runs (Miseq 2×300 bases) spiked with PhiX genomic DAN to ensure high enough sequence diversity. Fourteen environmental factors (Fig. 2B) were collected from TerraClimate [47] for each sampling month (Table. S3). The TerraClimate database has a monthly temporal resolution and approximately 4 km (1/24th degree) spatial resolution [47].

### Amplicon sequencing data analysis

Amplicon sequencing data was processed in Mothur (version 1.42.3) [48, 49] as described in Almario *et al*. [29]. Single-end reads were combined to make paired-end reads (make.contigs command), and paired reads with less than five bases overlap between the forward and reverse reads were removed. Only 100-600 bases long reads were kept (screen.seqs). Chimeric sequences were detected and removed using Vsearch [50] in Mothur (chimera.vsearch, remove.seqs). Cutadapt 2.10 [51] was used to trim primer sequences from 16S rRNA and 18S reads. For fungal reads, we used ITSx 1.1b [52] to trim reads to only the ITS2 region. Sequences were clustered into Operational Taxonomic Units (OTUs) at 97% similarity threshold (cluster, dgc method), and abundance filtering was applied to retain OTUs with more than 50 reads (split.abund) and OTU tables were generated (make.shared). OTUs were taxonomically classified (classify.otu) based on the Silva database [53] (version 138.1) for bacterial 16S rRNA data, the UNITE_public database [54] (version 02_02_2019) for fungal ITS2, and the Pr2 [55] (version 4.12.0) for eukaryotic 18S rRNA. The PhiX genome was included in each of the databases to improve the detection of remaining PhiX reads. OTUs classified as chloroplast, mitochondria, Arabidopsis, Embryophyceae, unknown, and PhiX were removed (remove.lineage).

### Diversity and mul9variate analysis

OTU tables (bacteria, fungi and non-fungal eukaryotes) were modified by removing samples with less than 50 reads. OTU abundance tables were used to calculate Shannon’s H−diversity index (estimate richness function in Phyloseq [56] R package) to estimate alpha−diversity. To calculate between-sample diversity, relative abundance OTU tables were computed and transformed (log10 (x + 1)), to calculate Bray-Curtis dissimilarities used for nonmetric multidimensional scaling ordination (NMDS, ‘ordinate’ function, Phyloseq [56] R package). A PERMANOVA analysis on Bray-Curtis dissimilarities was performed to identify the main factors influencing the structure of the leaf microbiome (‘adonis2’ function, Vegan package [57], 10 000 permutations, *P <* 0.05, explanatory categorical variables: Compartment x Site x Season). To facilitate comparability, all quantitative environmental variables (e.g., Temperature and Precipitation) were z-transformed to have a mean of zero and a standard deviation of one. These data were correlated to the measured alpha-diversity (as mentioned above) of each compartment (‘cor.test’ function, spearman method, Stats [58] package, *P <* 0.05). Environmental data was then used in PERMANOVA analyses to assess the effect of each factor on Bray-Curtis dissimilarities. In detail, for each microbial group and compartment, 14 models were performed (‘adonis2’, Vegan package [57], 10 000 permutations, *P <* 0.05, explanatory categorical variables: one environmental factor). Unless otherwise stated, data normality was checked (‘shapiro.test’, Stats [58] package), and means were compared using the nonparametric multivariate test for multiple groups (‘dunnTest’ function, FSA [59] package, Benjamini-Hochberg *Padj <* 0.05) and two groups (‘wilcox.test’ function, stats [58] package, *P <* 0.05). All analyses were performed in R (version 4.1.2) [60].

### Linear Model Analysis

The association between independent variables (environmental factors) and the dependent variable (relative abundance of genera) was investigated using linear models. Original abundance OTU tables (samples with *>* 1 read) were aggregated at the taxonomic genus level (aggregate function in R). Rare genera (those with *<* 50 reads) were excluded, and the table was converted to relative abundance. Additionally, highly correlated environmental factors were identified, and one factor, actual evapotranspiration, was removed from the analysis. A z-transformed environmental table was utilized for consistency. Linear models (‘lm’ function, stats [58] package) were executed per compartment per genus using the formula:

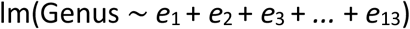

where ‘*e*’ denotes environmental factors. The resulting models were employed to identify the most influential environmental factors (ols−step−best−subset function, olsrr package [61]). . Models with the lowest estimated prediction error (msep parameter) were selected. To estimate the coefficient values of selected factors per genus, a generalized linear model was performed using the formula:

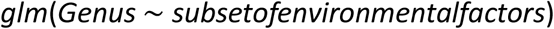

The significance of each factor individually (by dropping it from the model) was assessed (‘drop1’ function, ‘Chisq’ test, lme4 [62] package). The obtained *P*-values from the chi-squared tests were adjusted for false discovery rate (‘p.adjust’ function method=‘fdr’, stats [58] package, P *<* 0.05). Coefficient values (of selected environmental factors, demonstrating their strength in predicting the relative abundance of microbes, were utilized to subset some of the genera that exhibited differential effects between both compartments for further visualization. To do this, absolute values of coefficient values of environmental factors per genus were averaged, and the top overlapping genera between epiphytic and endophytic compartments were selected. All analyses were performed in R (version 4.1.2) [60].

### Microbial network calculations and properties

Bacteria, fungi, and non-fungal eukaryotes OTU abundance tables were merged and used for correlation calculation using the SparCC algorithm [63], which relies on Aitchison’s logratio analysis and is designed to deal with compositional data with high sparsity. OTU tables were filtered to OTUs in at least 5 samples with ≥ 10 reads per OTU per time point per compartment. The filtered OTU tables (OTU raw abundances) were used to calculate SparCC correlation scores (with default parameters) in FastSpar platform [64]. Pseudo p-values were inferred from 1000 bootstraps. Only correlations with *P* ≤ 0.001 and absolute correlation *>* 0 were kept for further analyses. Cytoscape (version 3.7.1) [65] was used for network visualization. A “Cohesion” metric [32] was calculated to quantify the connectivity of each network. For each sample (j), a positive and a negative cohesion metric (equation1) were calculated by multiplying OTUs relative abundances to the average of the OTU’s positive or negative correlations.

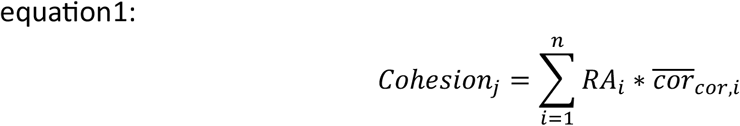

Where *RA*_*i*_ is relative abundance of *OTU*_*i*_ in sample j and 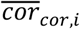 is average of significant positive (range from 0 to +1) or negative (range from -1 to 0) correlations for *OTU*_*i*_. Total cohesion per sample is then measured by the sum of the positive and negative cohesions. Total cohesion is correlated with environmental factors using Spearman correlation (cor function in R).

## Availability of data and material

Sequencing data are available under NCBI Bioproject PRJNA961058. OTU tables and scripts are available here https://gitlab.nfdi4plants.de/maryam mahmoudi/AbioticAraMicrobe

## Author Contributions

MM, KN and EK conceived and devised the study. MM, JA and KL performed the experiments. MM and JA analyzed the data. MM, JA, KN and EK contributed to writing and preparation of the manuscript. All authors read and approved the final manuscript.

## Acknowledgements

We thank KemenLabSamplingTeam for organizing and participating in several sampling trips and Elke Klenk for helping in MiSeq sequencing preparation. This project has been funded by the European Research Council (ERC) under the DeCoCt research program (grant agreement: ERC-2018-COG 820124), the Cluster of Excellence “Controlling Microbes to Fight Infections” (CMFI; Exc 2124) and the SPP 2125 DECRyPT program from the DFG.

## Supplementary figures

**Supplementary figure 1.**
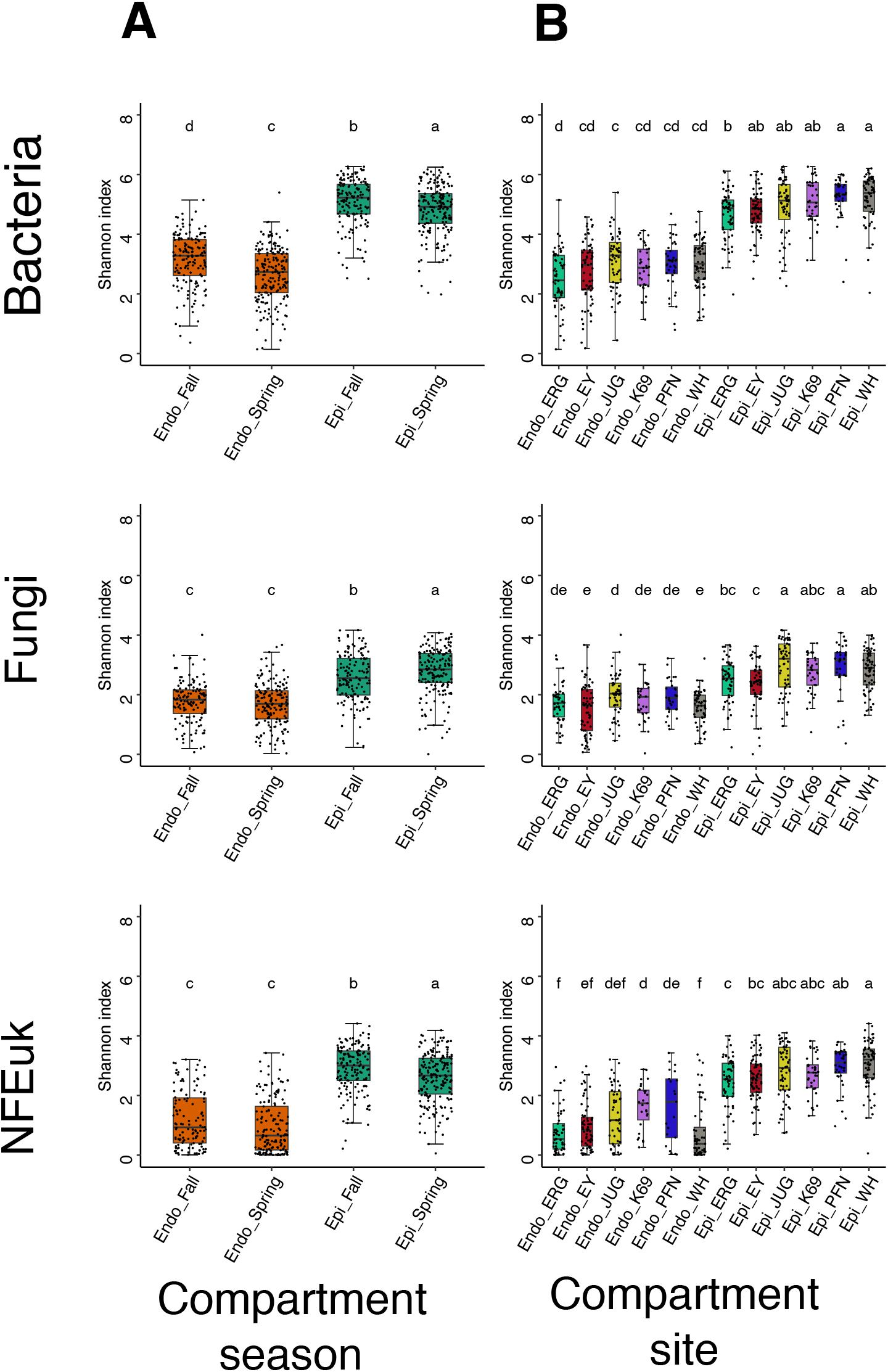
Within-sample diversity of leaf compartments across sampling sites and sampling seasons. Alpha diversity of epiphytic and endophytic samples across spring and fall samples **(A)** and sampling sites **(B)** in bacteria, fungi, and non-fungal eukaryotic communities. The box plots display individual samples as dots. Different letters indicate significant group differences (Dunn test, *P <* 0.05).

**Supplementary figure 2.**
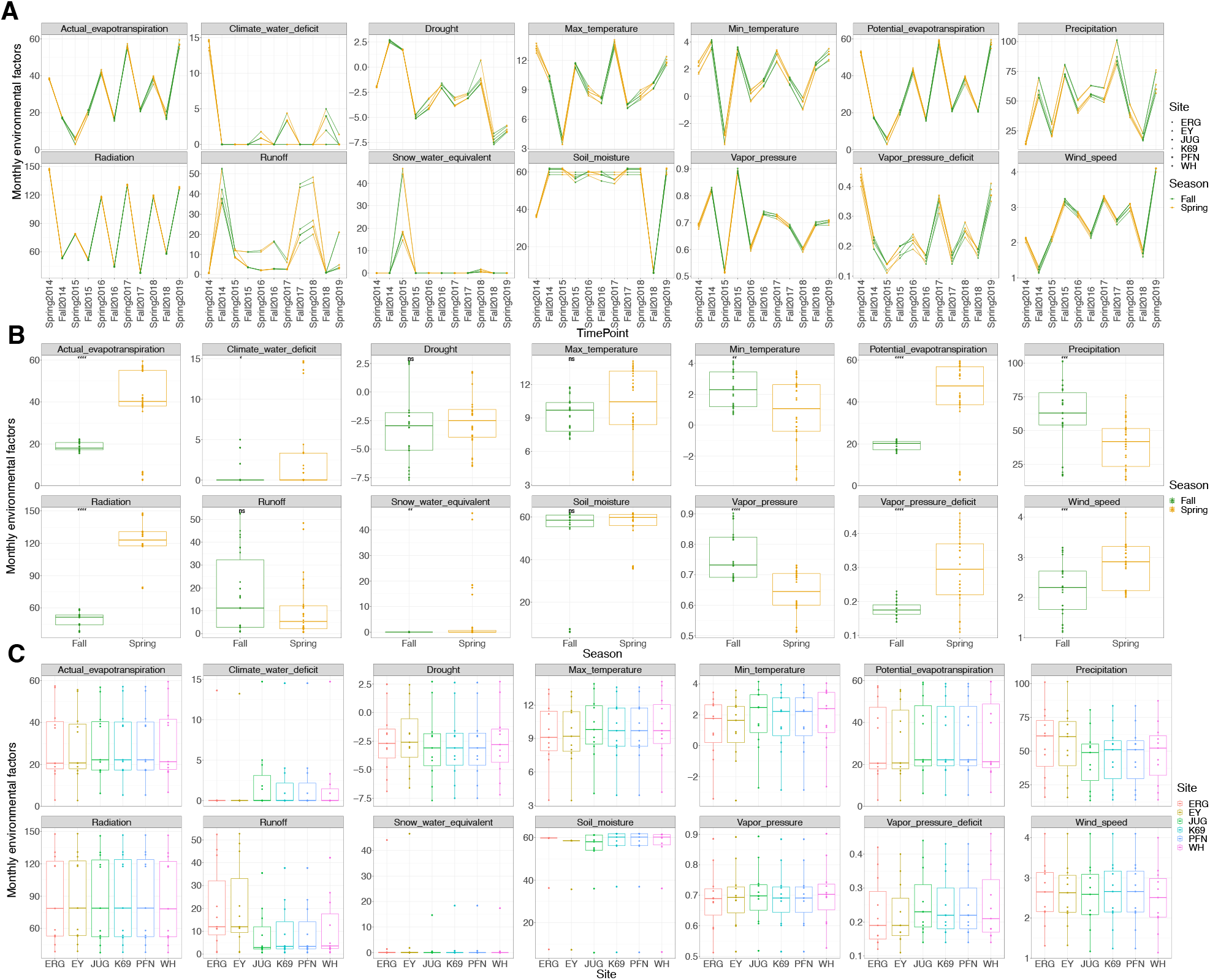
Changes in environmental variables over sampling time points and sampling sites. **(A**) Lines show the average values for each environmental factor for each sampling month and site. **(B)** Spring vs fall differences for each environmental factor **(C)** Differences between sites for each environmental factor. Asterisks indicate significant differences based on Wilcoxon’s test: n.s. (*P >* 0.05), * (*P* ≤ 0.05), ** (*P* ≤ 0.01), *** (*P* ≤ 0.001), and **** (*P* ≤ 0.0001).

**Supplementary figure 3.**
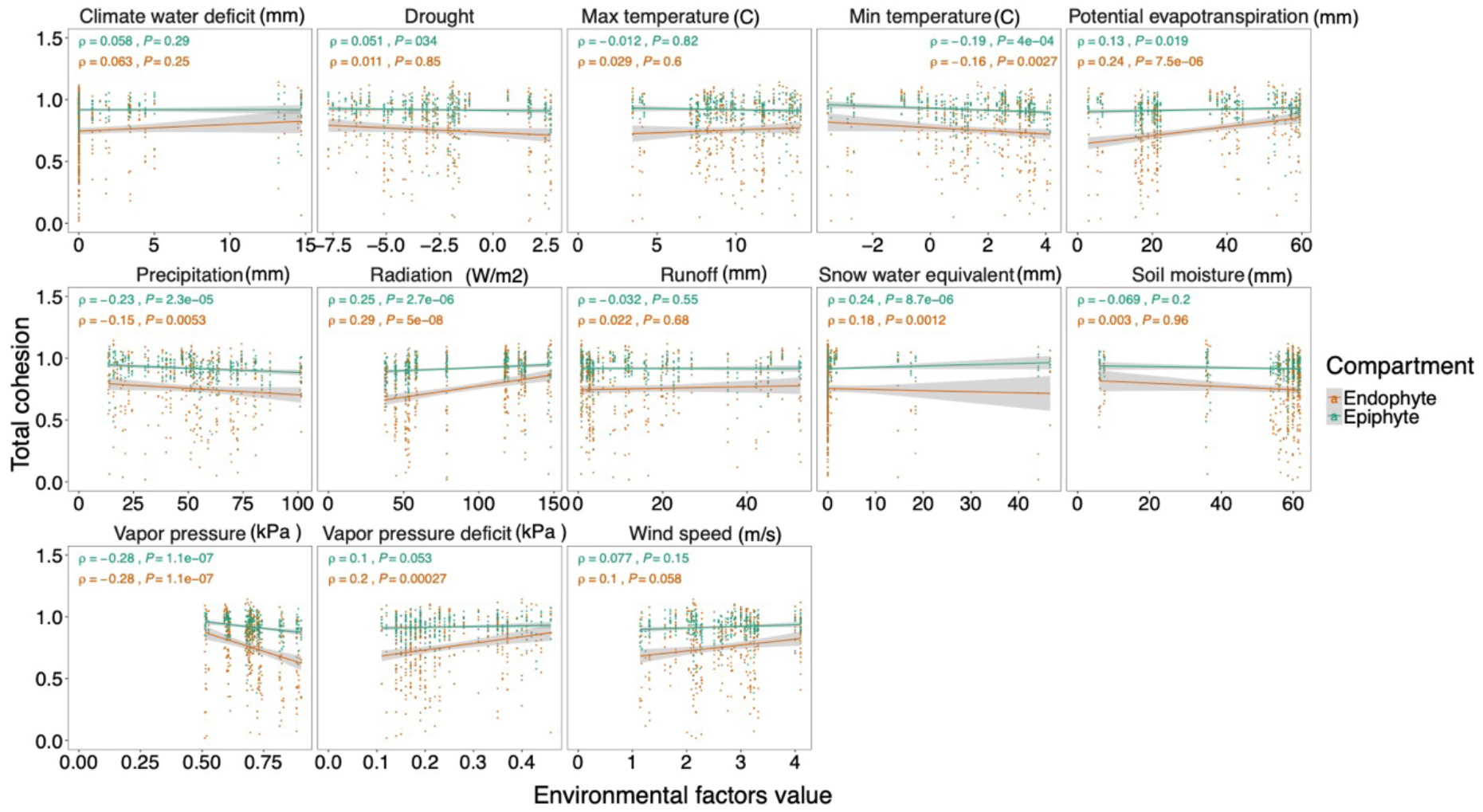
Relationship between total cohesion and environmental factors. Each plot shows a linear regression model fit to the data to show the association of total cohesion and environmental factors, across compartments. Samples are represented by dots and colored by compartments. The grey lines indicate 95% confidence intervals, and the Spearman correlation coefficient *ρ* and actual *P*-values are shown.

**Supplementary tables are in https://gitlab.nfdi4plants.de/maryammahmoudi/AbioticAraMicrobe**

**Table S1. Experimental set-up with sampling locations and sampling time points**.

Numbers indicate the number of samples collected from epiphytic and endophytic samples at different time points (season and year). In addition, the exact date of each sampling event and the coordinates of the sampling locations are given.

**Table S2. Fourteen environmental factors (Fig. 1B) used in this study**.

The environmental variables were obtained from TerraClimate [47], a database with monthly temporal resolution and approximately 4 km spatial resolution.

**Table S3. PERMANOVA results indicating the impact of individual environmental factors on microbial communities within each leaf compartment**.

**Table S4. Results of linear modeling association of environmental factors and microbial genera**.

**Table S5. Pearson correlation analysis results depicting the relationship between the number of nodes in microbial networks and the environmental factors**.

**Table S6. Primers and blocking oligos used in this study**.

